# Volumetric fluorescence microscopy-based quantitative comparison of murine tissue clearing using CUBIC protocols

**DOI:** 10.64898/2026.03.06.709534

**Authors:** Roland Pohlmeyer, Sergiy V. Avilov, Wolf Heusermann, Dagmar Diekhoff, Oliver Biehlmaier

## Abstract

A wide variety of protocols have been proposed for optical clearing of tissues, whole-mount organs, and other bulky specimens to enable their volumetric fluorescence imaging. However, quantitative comparisons of tissue clearing protocols that take into account the fluorescence of the final specimens remain rare. Here, we propose a volumetric fluorescence image–based workflow for evaluating tissue clearing and fluorescence staining protocols. The workflow calculates depth-dependent fluorescence attenuation coefficients using data from entire 3D images, thereby avoiding spatial sampling bias and eliminating reliance on simple readouts, such as light transmittance, to predict fluorescence image quality. By combining autofluorescence signal with the signal from a specific fluorescence label, we independently evaluated transparency and the quality of fluorescence staining in cleared specimens. Using the proposed workflow, we systematically compared clearing and staining performance of three CUBIC-based protocols in murine liver, kidney, spleen, thymus, and intestine, and revealed differences in final fluorescence image quality across protocol–organ combinations.

## Introduction

A detailed understanding of how tissues and cells are arranged in three dimensions is essential for many areas of biology. Ideally, this would be achieved by imaging intact specimens in their entirety; however, most biological tissues cannot be directly visualized in depth because they are naturally opaque due to their heterogeneous composition - organelles, cytoplasm, extracellular matrix, chromatin, and other structures, all possessing distinct refractive indices (RI). These refractive index mismatches produce scattering and reflection of light as it passes through biological tissue, restricting the depth at which sharp images can be acquired [1]. To overcome this limitation, diverse methods were developed since over a century [2], aiming to equalize the refractive index throughout the specimen, typically by infiltrating it with high-RI reagents and by removing strongly light-scattering components such as lipids, and, in some cases (for example CUBIC reagents [3]), by removing also light-absorbing substances (i.e. pigments). Over last decades, wide variety of these methods, named optical tissue clearing, emerged (for example, [4] [5–8] [3,9,10]; their comprehensive review is beyond the scope of present Introduction). Tissue clearing protocols are roughly divided into four groups: organic solvent-based (hydrophobic), simple immersion, hyperhydration and hydrogel-embedding [1,11], which differ substantially in the level of transparency achieved, their practicality, compatibility with various fluorophore types and other parameters [11]. Additional parameters should be considered for the protocols used in multi-user core facility environment. Ideally, these protocols have to be suitable for various tissues and multiple types of fluorescent labels, including fluorescent proteins; the reagents should not be highly hazardous and should be compatible with dipping objectives of the microscopes used for volumetric imaging at the core facility. Considering all these, in our core facilities we have implemented the tissue clearing protocols based on CUBIC reagents, adapted from the original publications [3,9,10]. Besides the above-mentioned advantages, CUBIC reagents remove heme-derived pigments [10] which absorb visible light, thus significantly contributing to fluorescence attenuation in many vertebrate organs (*e.g.* liver and spleen). Performance of any given tissue clearing protocol depends on the type of tissue and the fluorophores to be imaged. For choosing an appropriate tissue clearing strategy, systematic comparisons (quantitative, if possible) of various protocols are very instrumental. A number of comparisons of tissue clearing protocols were published. Orlich et Kieffer systematically compared ten tissue clearing protocols qualitatively, on the basis of visual appearance of the cleared specimens [12]. As a quantitative measure of clearing efficiency, the most obvious and simple is light transmittance, used for example in [13], and also suitable for high throughput ([10,14]). However, typical goal of clearing a whole organ or tissue is acquisition of its 3D fluorescence images, whose quality depends not only on transparency, but also on the quality of fluorescence labelling: how well the label penetrated the tissue and bound its specific target, how well its fluorescence is preserved, etc. Fluorescence of the cleared tissues was evaluated in few publications (*e.g.* [15], [13]). For example in [16,17], signal to noise ratio (SNR) of fluorescence images was calculated, however, only few individual intensity profiles (two-dimensional plots of intensity as a function of a spatial coordinate), drawn in few selected optical sections were used. In heterogeneous specimens (such as a whole-mount organs or tissues), such values are sensitive to exact positions of these line profiles within 3D images, thus introducing spatial sampling bias. In [18], signal to background ratio was calculated for several 10 μm×10 μm regions per specimen. In comparison to line profiles, this reduced sampling bias, but has not completely eliminated it. In the cited work, intensity profiles of z-projections were plotted to visualize fluorescence attenuation across tissue specimens, however, without any quantitative parameter of this depth-dependent attenuation.

Here, to avoid the spatial sampling bias, we use the averaged fluorescence intensity profile from the entire imaged volume of the cleared specimen, which we named “global intensity profile”. From these profiles obtained in the autofluorescence and the specific fluorescence channels, we calculated depth-dependent fluorescence attenuation coefficients, which served as integral quality metrics of cleared fluorescently labelled specimens. Namely, attenuation of autofluorescence was used to estimate transparency, while attenuation of a specific fluorescence label permitted to infer the quality of fluorescence staining. Generally, autofluorescence is considered undesirable in fluorescence microscopy (e.g. [19]), since it contributes to background signal which does not originate from specific fluorescent markers of interest. However, if a color channel is “reserved” for autofluorescence (no specific fluorescent label is detected in that channel) then autofluorescence may be useful to visualize tissue context. For instance, it was used for imaging of whole-mount human tissues [20]; applications of autofluorescence in microscopy are reviewed in [21]. For our purpose, most important is that autofluorescence does not depend on labelling efficiency nor on label penetration into the tissue. Therefore depth-dependent attenuation of autofluorescence in tissue should only depend on its transparency (assuming no systematic gradient of autofluorescent substances), thus permitting to estimate transparency without contribution of any label distribution. On the other hand, specific staining (with an antibody, a DNA-binding dye etc) is a time-consuming and critical step of a whole-mount specimens preparation, because the reagent has to penetrate thick tissue and (in ideal case) to stain uniformly its specific molecular target. Inefficient penetration or poor binding to specific target may hamper extraction of useful information from the images, even for perfectly transparent specimens. Therefore, quality of exogenous labelling of whole-mount samples needs characterization as well. In general, depth-dependent attenuation of specific fluorescence in the tissue may depend on both: 1) penetration of the dye into the tissue, and 2) tissue transparency. In the present study, attenuation of the specific fluorescence permitted us to infer specific labelling-related processes, since transparency was independently estimated from attenuation of autofluorescence. As a specific fluorescent staining, we have used DNA-binding dye propidium iodide (PI). Furthermore, we have systematically applied the proposed workflow to compare quality of clearing and PI fluorescence staining of murine liver, kidney, spleen, thymus and intestine by 3 CUBIC-based tissue clearing protocols (summarized on Fig 1).

**Fig 1.**
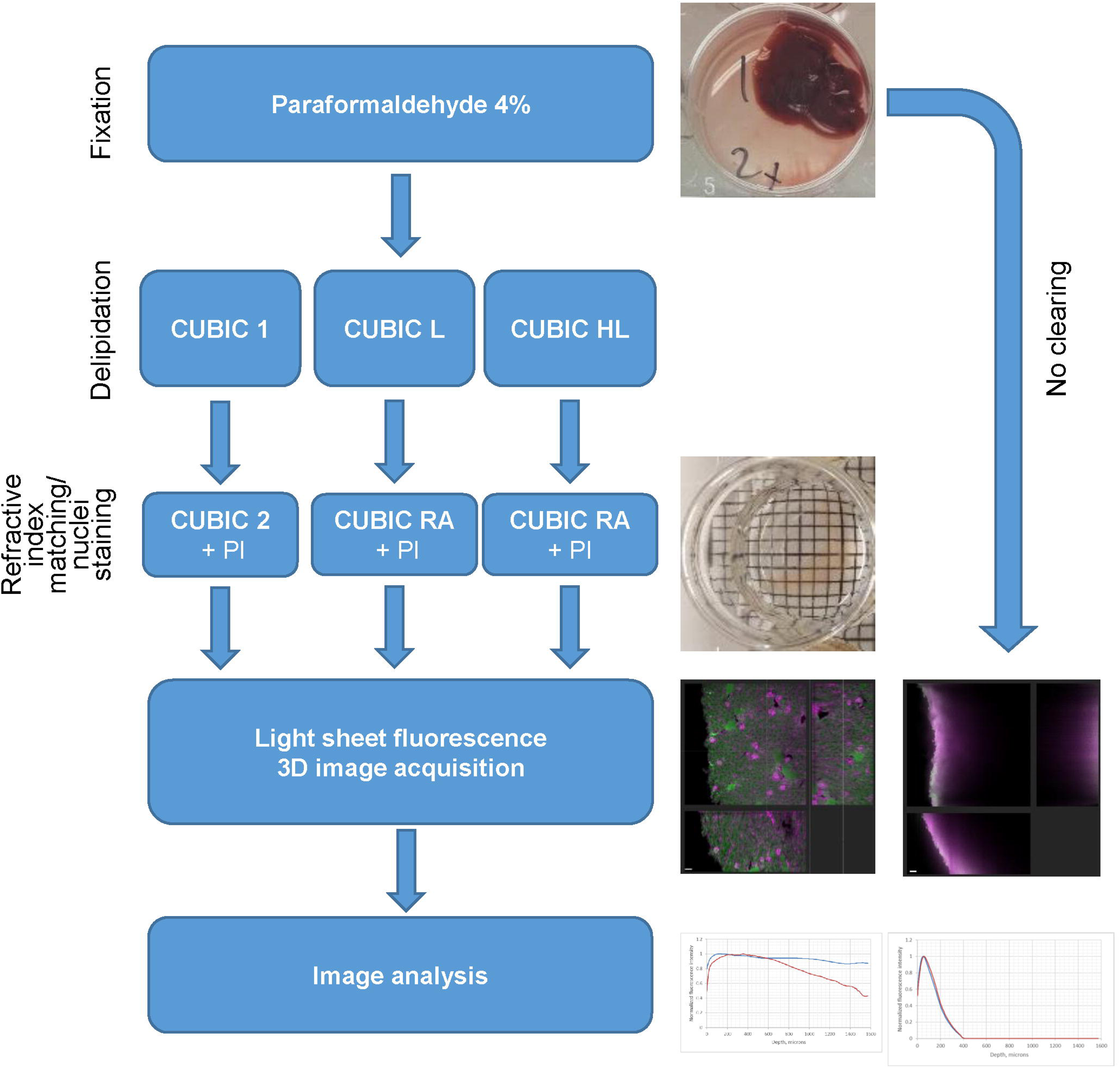
Conceptual diagram of the specimen preparation, image acquisition and analysis.

## Materials and Methods

### Reagents and materials

Reagents, materials and preparation of working solutions are described in [22].

### Animals and tissues

All mice were raised in the animal facility of the Max Planck Institute of Immunobiology and Epigenetics under specific pathogen-free conditions and killed according to the German animal welfare rights. All animal experiments were performed in accordance with the relevant guidelines and regulations, approved by the review committee of the Max Planck Institute of Immunobiology and Epigenetics and the Regierungspräsidium Freiburg, Germany. Murine organs were obtained from adult B6 wild type mice, strains C57BL76J and C57BL/6NTac. The thymus for the supplementary images with fluorescent proteins were taken from an adult mouse from the line Tg(pFoxn1-Turquoise-mCherry)Tbo/Mpie. In this line, nuclei of thymic epithelial cells are red fluorescent, whereas their cellular membranes are marked by cyan fluorescence. A detailed description can be found in [23].

### Optical tissue clearing and fluorescence labelling

Optical tissue clearing and fluorescent labelling protocols are described in detail in [22]. In comparison to the original protocols [10], the following modifications were made. 1) The refractive index of CUBIC RA prepared according to the original publication [10] is 1.52. To adjust it to 1.45 (the optimal value for Lightsheet.Z1 optics), we diluted CUBIC RA with water until the target RI value was reached (controlled by a refractometer). 2) Propidium iodide was added directly to refractive index matching solutions (CUBIC 2 or CUBIC RA), so nuclei staining step was combined with the refractive index matching.

### Light sheet fluorescence microscopy (LSFM) image acquisition

LSFM images were acquired on Lightsheet.Z1 system (Zeiss), using illumination objective LSMFM 5x/0.1, detection objective EC Plan Neofluar 5x/0.16, PCO edge CMOS cameras (Hamamatsu) at 200 ms exposure, using one-side illumination. Spatial sampling was 1.21 microns in *xy* and 5.89 microns in z. Fluorescence emission was spectrally separated into two channels using dual-view emission splitters equipped with a dichroic mirror and two band pass emission filters, directing the signals onto two synchronized cameras. Namely, autofluorescence and PI signals were excited with 488 nm and 561 nm lasers respectively, and separated using the emission splitter with SBS LP 560 dichroic and collected with BP 505-545 emission filter (“GFP” channel) and LP 585 emission filter (“RFP” channel). mTurquoise and mCherry signals of the thymi expressing these FPs (S2 Fig) were excited with 445 nm and 561 nm lasers, respectively, and their emission was splitted using the emission beam splitter equipped with SBS LP 510 dichroic, BP 460-500 emission filter for mTurquoise, and BP 575-615 emission filter for mCherry. The specimen illumination, emission detection and positions of the intensity profiles are shown on the Fig 2. The excitation light sheet propagated along the x coordinate from left side only, such that the attenuation of the emitted fluorescence along x axis represented the decay of excitation light with the length of its path in the tissue. The emitted fluorescence was “viewed” by the camera from z direction.

**Fig. 2.**
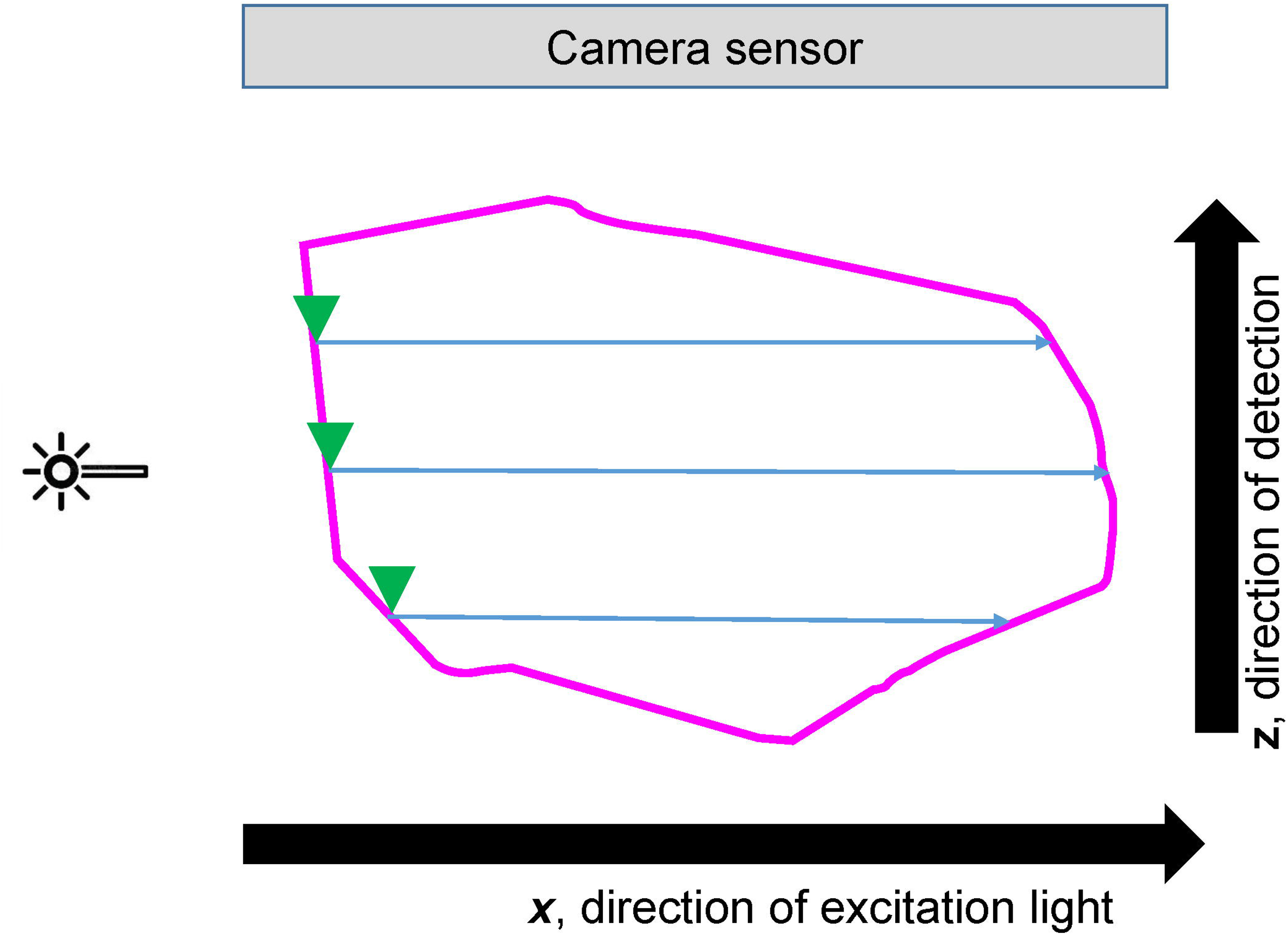
Scheme of beam paths and intensity profiles orientation (corresponding to the top view in Lightsheet.Z1 instrument). Single *xz* plane is shown for clarity. Excitation light propagates in *x* direction. Bold magenta line represents *xz* section of the surface of the tissue fragment. Blue arrow lines represent individual intensity profiles; for clarity, only few profiles are shown. Green arrowheads indicate positions where the depth was set to zero for the shown intensity profiles. Emitted fluorescence is detected along *z* axis. For the analysis, all intensity profiles within a 3D tissue fragment were averaged.

### Image analysis

Two-channel 3D LSFM images were subject to a multi-step image analysis workflow combining GUI-based and scripted components. Raw fluorescence images were segmented on the basis of autofluorescence channel with Imaris 9.2 software (Andor), *Surfaces* tool (icmx file containing the exact segmentation parameters is available on request). Using the obtained *Surfaces,* the signals in the both “GFP” and “RFP” channels were masked to set values outside the segmented volume to zero, and then exported as tiff file series. Further, a custom Python script (available on request) was applied to calculate the global intensity profiles in the following way. Since the tissue fragments did not occupy the entire imaged 3D volumes, the absolute x-coordinate for each intensity profile was adjusted individually to align the tissue surface to *x*= 0 (Fig 2) (*i.e*., the position at which the excitation beam entered the tissue was defined as *x*=0). All individual intensity profiles (*x*) obtained at each *y*-position within the tissue in a given *xy* optical slice were averaged, yielding one profile per slice. These slice-averaged profiles were then averaged across the entire *z*-stack to obtain a single intensity profile representing the entire volume of the sample, further referred to as “global intensity profile”. These global intensity profiles were used to calculate fluorescence attenuation coefficients as following. We linearized the exponential decay model *I*(*x*) ≈ *I*_0_*e*^−µ*x*^by taking *y* = *ln*(*I*/*I*_0_) (where *I* is the fluorescence intensity at the depth in tissue equal to *x,* and *I_0_* is the intensity at the surface of the specimen), and fitted a robust linear slope using the Theil–Sen (median-of-pairwise- slopes) estimator; the attenuation coefficient was reported as *μ* = −*s*^ [24]. Point estimates of *μ* were accompanied by 95% confidence intervals obtained by nonparametric bootstrap resampling (1000 resamples) of the measured depth points. The Theil–Sen estimator and bootstrap intervals were computed using scipy.stats.theilslopes and a custom resampling Python script (available on request).

### Statistical analysis

All specimens were prepared in triplicates, except the non-cleared liver (Fig 8) and the thymi containing mTurquoise and mCherry (S2 Fig). Median values of fluorescence attenuation coefficients were calculated for each condition (Fig 9).

## Results

### The workflow for quantitative metric of clearing efficiency

The experimental workflow is summarized in Fig 1. The specimens (fragments of murine organs) were cleared according to one of the indicated protocols; their 3D LSFM datasets were acquired in the “GFP” channel, dedicated to autofluorescence, and in the “RFP” channel used for fluorescence of PI. These datasets were analyzed to quantify depth-dependent fluorescence attenuation (*Materials and Methods*). Briefly, tissue volumes were segmented based on autofluorescence; within these segmented volumes, fluorescence intensity profiles were averaged across lateral positions and then through axial slices to generate a single global intensity profile per specimen (Fig 2). These profiles were then used to calculate fluorescence attenuation coefficients.

### Organization of results

The results are presented in the following way: Figs 3-7 contain orthogonal views of representative LSFM 3D images of the cleared tissues, and the corresponding global intensity profiles for these specimens. Fig 8 shows orthogonal views and global intensity profiles of a non-cleared liver fragment. Fig 9 summarizes fluorescence attenuation coefficients for all analyzed organ/clearing protocol combinations. LSFM images of thymus fragments expressing mTurquoise- and mCherry-tagged proteins are in the S2 Fig. Global intensity profiles for all replicates of LSFM datasets for autofluorescence and PI signals, respectively, are shown on S3 and S4 Figs.

**Fig 3.**
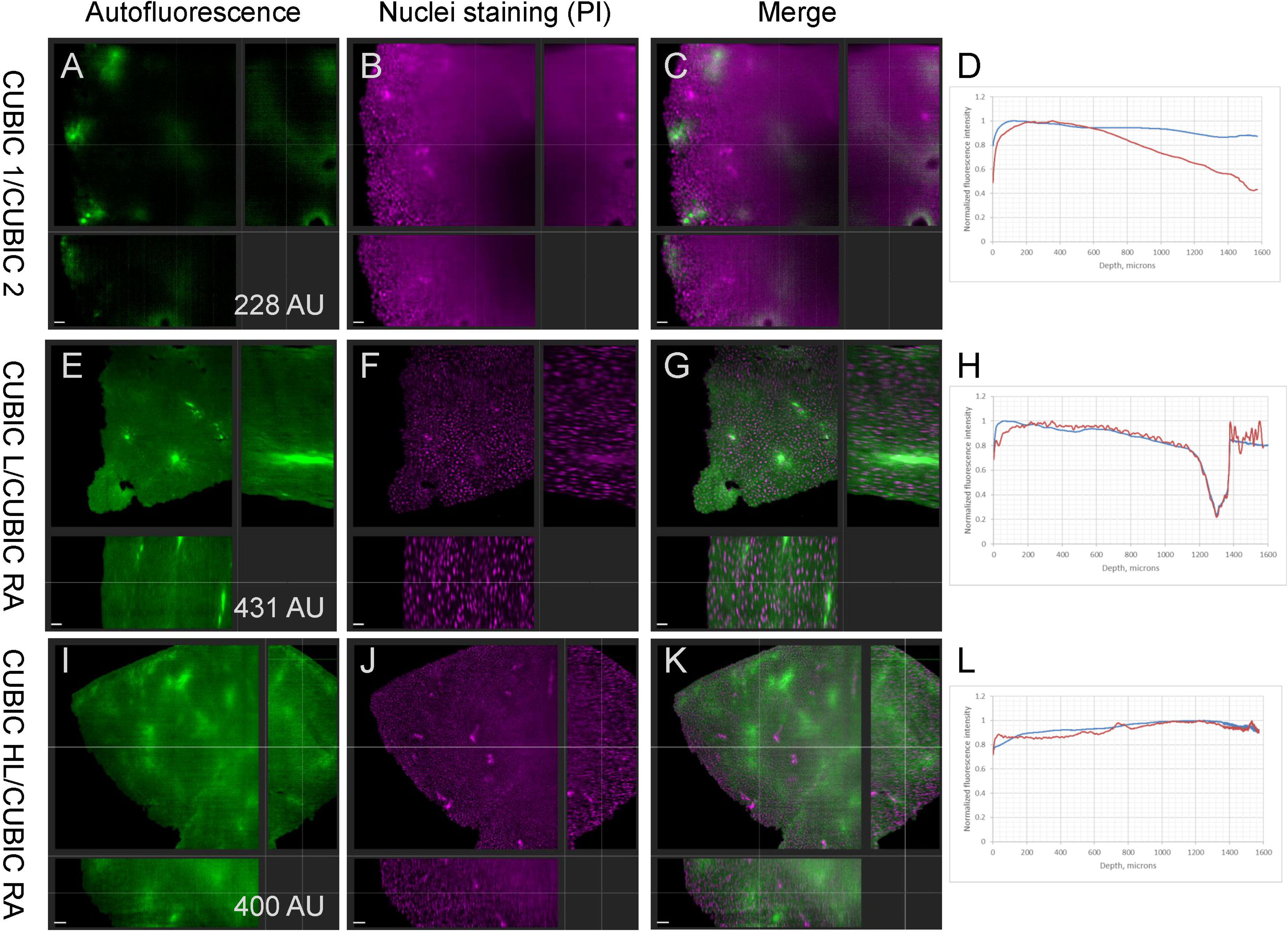
Ortho projections of 3D LSFM datasets of liver tissue cleared with CUBIC 1/CUBIC 2 (A-C), CUBIC L/CUBIC RA (E-G) and CUBIC HL/CUBIC RA (I-K) protocols. For visualization purposes, brightness of the “GFP” channel is individually adjusted in each panel. Mean absolute intensities in “GFP” channel are marked on the panels A, E and I. Scale bar 100 microns. D, h and l: global intensity profiles extracted from the respective 3D LSFM datasets: profile of autofluorescence channel (blue) and profile of PI channel (red).

**Suppl. figure 1. Visual appearance of representative liver, intestine, kidney, and spleen tissue through the steps of the tissue clearing protocols used in the present paper**.

**Suppl. figure 2. Ortho projections of 3D LSFM datasets of thymus tissue expressing mTurquoise and mCherry, cleared with CUBIC 1/CUBIC 2 (A-C) and CUBIC L/CUBIC RA (E-G) protocols.** Scale bar 100 microns.

**Suppl. figure 3. Global intensity profiles in the autofluorescence channel for all analysed datasets. Suppl. figure 4. Global intensity profiles in the PI staining channel for all analysed datasets.**

### Comparison between the tissue clearing protocols

All the three tested protocols permitted us to clear, at least partially, the tested organs (Figs 3-7), enabling better light penetration than in the non-treated tissues (Fig 8). The fluorescence attenuation coefficients (Fig 9) of all cleared specimens were substantially lower than those of the non-cleared tissue (9.9 mm^−1^ for autofluorescence and 9.7 mm^−1^ for nuclei staining, not shown). However, clearing efficiency varied to some extent between organs and protocols. Overall, CUBIC L/CUBIC RA protocol yielded rather bright absolute autofluorescence, good transparency and uniform PI staining (Figs 3E, 4E, 5E, 6E and 7E), also indicated by low attenuation coefficients for both autofluorescence and PI fluorescence (Fig 9). CUBIC 1/CUBIC 2-treated specimens had dimmer absolute autofluorescence than the samples treated with other protocols (Figs 3A, 4A, 5A, 6A and 7A). CUBIC 1/CUBIC 2 protocol was less efficient than others for thymi and spleens, as indicated by rather high attenuation coefficients in the autofluorescence channel (Fig 9A). Finally, CUBIC HL/CUBIC RA permitted efficient clearing of all tissues (autofluorescence attenuation coefficients < 0.6 mm^−1^, Fig 9A), except spleens (attenuation coefficients for individual specimens ranging from 0.2 to 4 mm^−1^). CUBIC HL/CUBIC RA was the only protocol that yielded a low (nearly zero) and reproducible attenuation coefficient for thymi. However, treatment with CUBIC HL reagent (having pH>12), resulted in apparent dissolution of some of the specimens after few days of incubation. Therefore, to preserve their mechanical integrity, it was necessary to inspect specimens in CUBIC HL visually every day and to terminate CUBIC HL incubation as soon as they became apparently transparent.

**Fig 4.**
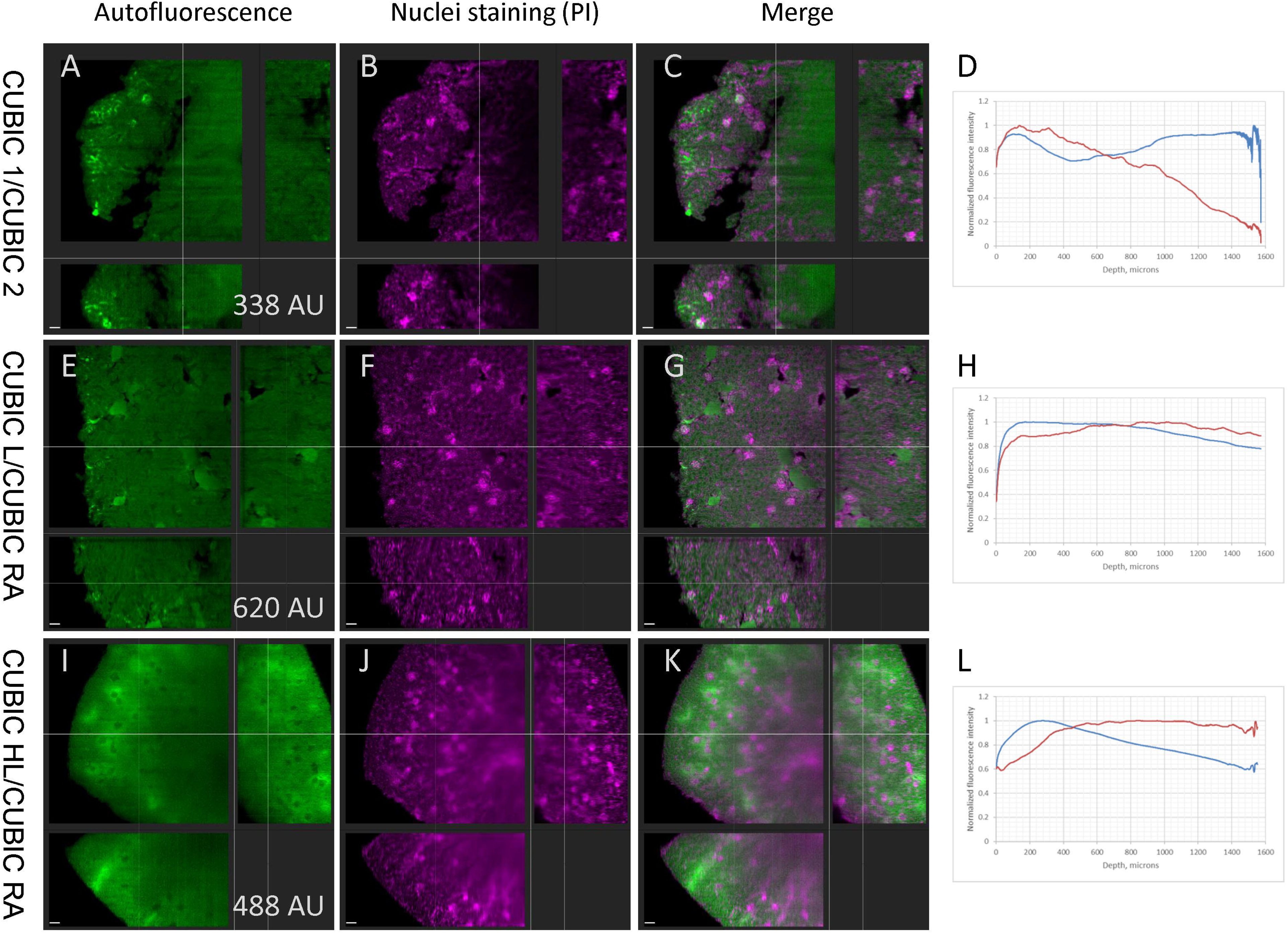
Ortho projections of 3D LSFM datasets of kidney tissue cleared with CUBIC 1/CUBIC 2 (A-C), CUBIC L/CUBIC RA (E-G) and CUBIC HL/CUBIC RA (I-K) protocols. For visualization purposes, brightness of the “GFP” channel is individually adjusted in each panel. Mean absolute intensities in “GFP” channel are marked on the panels A, E and I. Scale bar 100 microns. D, h and l: global intensity profiles extracted from the respective 3D LSFM datasets: profile of autofluorescence channel (blue) and profile of PI channel (red).

**Fig 5.**
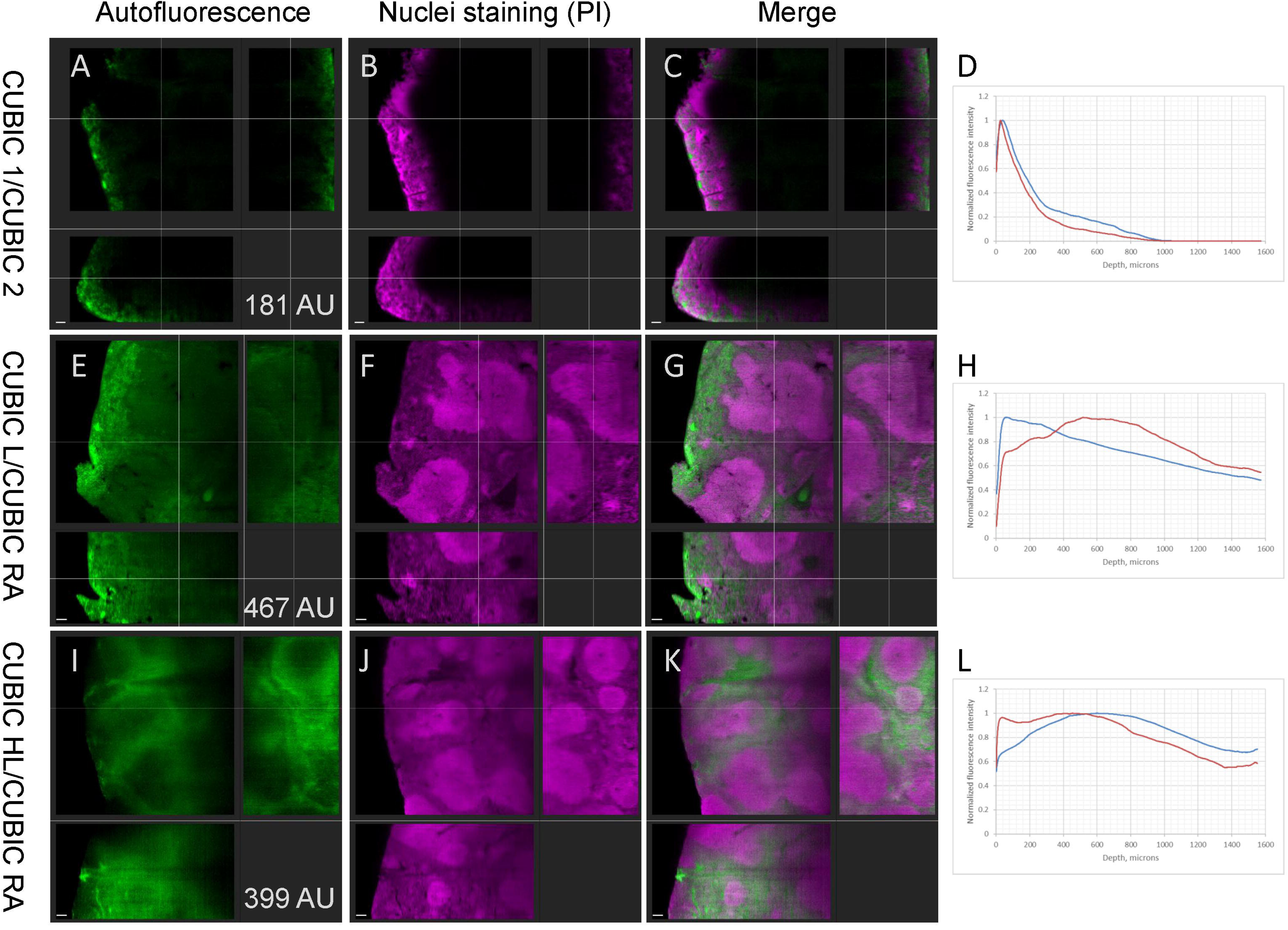
Ortho projections of 3D LSFM datasets of spleen tissue cleared with CUBIC 1/CUBIC 2 (A-C), CUBIC L/CUBIC RA (E-G) and CUBIC HL/CUBIC RA (I-K) protocols. For visualization purposes, brightness of the “GFP” channel is individually adjusted in each panel. Mean absolute intensities in “GFP” channel are marked on the panels A, E and I. Scale bar 100 microns. D, h and l: global intensity profiles extracted from the respective 3D LSFM datasets: profile of autofluorescence channel (blue) and profile of PI channel (red).

To reduce total sample preparation time, we performed nuclei staining simultaneously with RI matching (PI was dissolved directly in one of the RI matching solutions, CUBIC RA or CUBIC 2). Overall, this yielded efficient staining of the nuclei. However, in the CUBIC 1/CUBIC 2 specimens, PI fluorescence was more attenuated with the depth (median intensity attenuation coefficients of 0.67, 1.18, 2.28 and 1.87 mm^−1^, for liver, kidney, spleen and thymus, Fig 9B), than autofluorescence in the same conditions (respective intensity attenuation coefficients of 0.47, 0.12, 2.21, 0.38 mm^−1^, Fig 9A). Conversely, in CUBIC L/CUBIC RA samples, PI signal (intensity attenuation coefficients of 0.16, 0.21, 0.42 and 0.77 mm^−1^, for liver, kidney, spleen and thymus, Fig 9B) was roughly equally or less attenuated than that of autofluorescence in respective specimens (0.18, 0.64, 0.50 and 0.90 mm^−1^, Fig 9A).

### Comparison between organs

Quality of fluorescent images for each clearing protocol varied between organs. **Liver** and **kidney** were efficiently cleared with all the three protocols (Figs 3 and 4), as indicated by low autofluorescence attenuation coefficients (median values below 0.5 mm^−1^, Fig 9A). In the nuclei staining channel, CUBIC 1/CUBIC 2-treated liver and kidney looked more blurry (Fig 3F and Fig 4F) and had slightly higher attenuation coefficients, as discussed above. In the **spleens** treated with CUBIC 1/CUBIC 2, autofluorescence decayed steeply with depth (Figs 5A-D), with attenuation coefficient of 2.12 mm^−1^ (Fig 9A); in CUBIC HL/CUBIC RA spleens (Figs 5I-L), we obtained median attenuation coefficient of 1.38 mm^−1^ and a wide sample-to-sample variation (Fig 9A) for autofluorescence, indicating mediocre and poorly reproducible clearing. On the contrary, CUBIC L/CUBIC RA-treated spleens were cleared efficiently (Figs 5E-5H) and had rather low autofluorescence attenuation coefficient (median 0.5 mm^−1^, Fig 9A). **Thymi** treated with CUBIC HL/CUBIC RA had no obvious depth-dependent fluorescence attenuation in neither autofluorescence nor PI channels (Fig 6), with very low (nearly zero) attenuation coefficients in the both channels (Fig 9), while CUBIC 1/CUBIC 2- and CUBIC L/CUBIC RA-treated **thymi** showed widely varying autofluorescence attenuation coefficients (median values of 0.4 and 0.9 mm^−1^ respectively, Fig 9A). Incidentally, CUBIC L/CUBIC RA-treated thymi fragments looked inhomogeneous, especially in PI channel, where two distinct regions appeared in the images (Fig 6F): 1) those with bright nuclei on a dark background, and 2) the regions with uniform red fluorescence. The i**ntestine** fragments treated with CUBIC 1/CUBIC 2 and CUBIC L/CUBIC RA showed no apparent attenuation of neither autofluorescence nor PI fluorescence (Fig 7). Early during incubation in CUBIC HL, all the tested intestine fragments were dissolved, so it was not possible to acquire any LSFM images of CUBIC HL-cleared intestine fragment.

**Fig 6.**
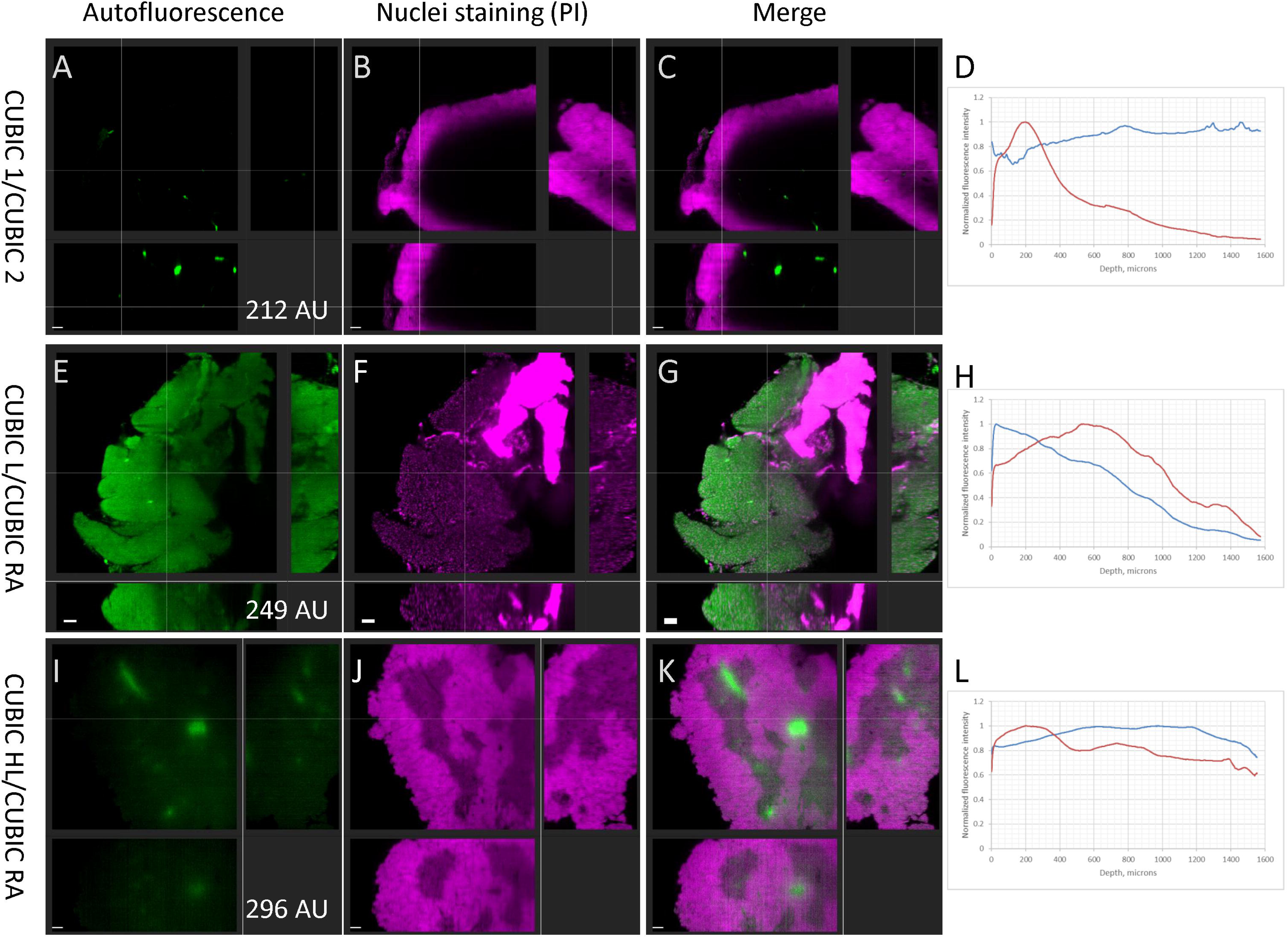
Ortho projections of 3D LSFM datasets of thymus tissue cleared with CUBIC 1/CUBIC 2 (A-C), CUBIC L/CUBIC RA (E-G) and CUBIC HL/CUBIC RA (I-K) protocols. For visualization purposes, brightness of the “GFP” channel is individually adjusted in each panel. Mean absolute intensities in “GFP” channel are marked on the panels A, E and I. Scale bar 100 microns; and global intensity profiles (D, H and I) extracted from the respective 3D LSFM datasets: the profile of autofluorescence channel (blue) and the profile of PI channel (red)

**Fig 7.**
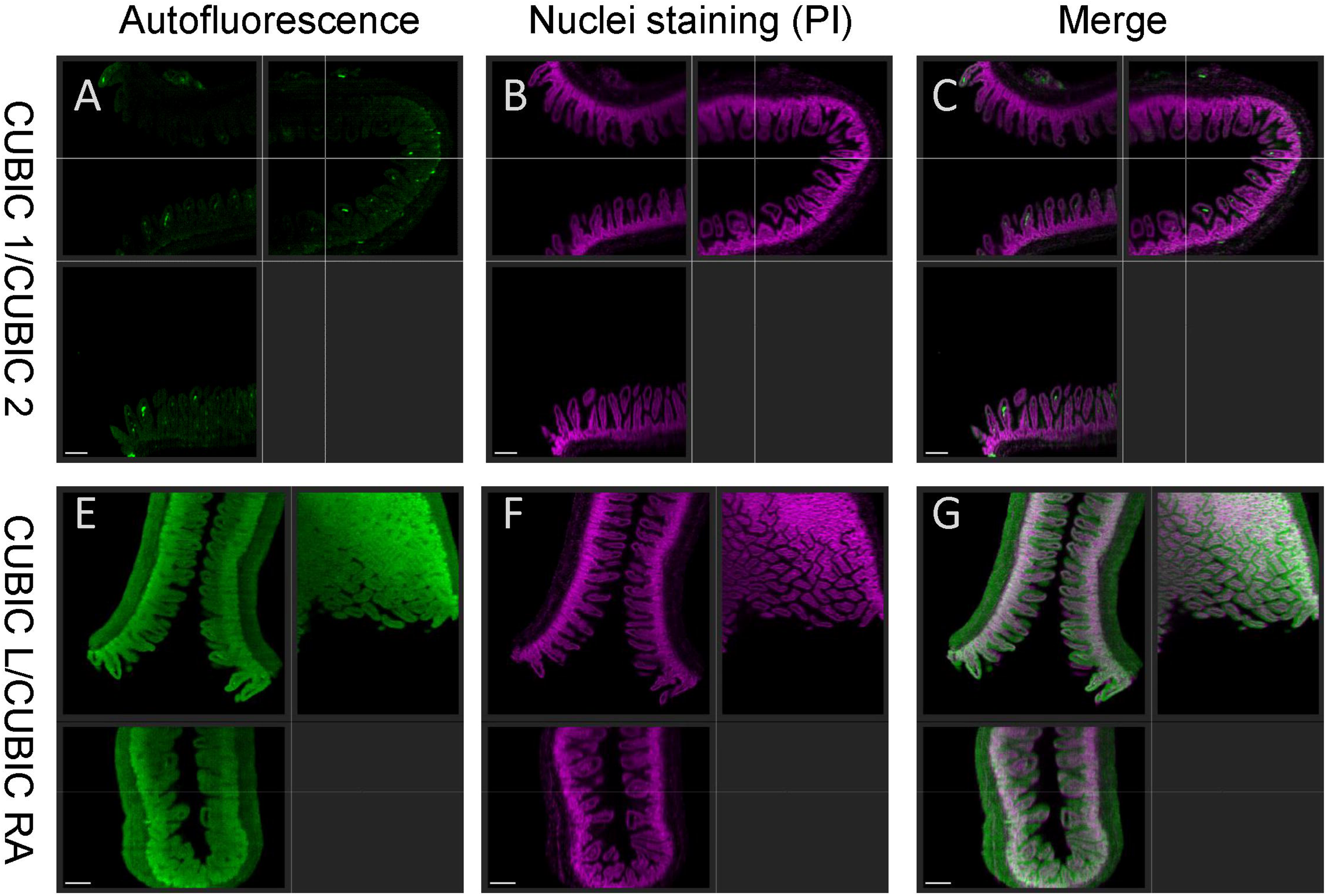
Ortho projections of 3D LSFM datasets of intestine tissue cleared with CUBIC 1/CUBIC 2 (A-C), and CUBIC L/CUBIC RA (E-G). Scale bar 100 microns.

**Fig 8.**
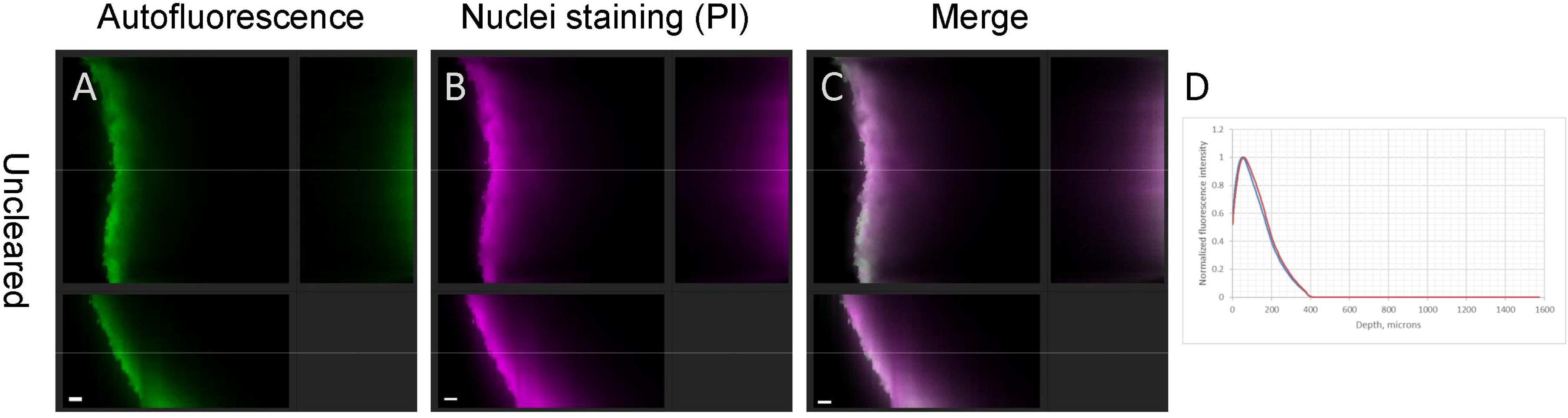
Ortho projections of 3D LSFM dataset of fixed non-cleared liver tissue (A-C) and the global intensity profiles extracted from the 3D LSFM dataset (D). The profile of autofluorescence channel (blue) and the profile of PI channel (red). Scale bar 100 microns.

**Fig 9.**
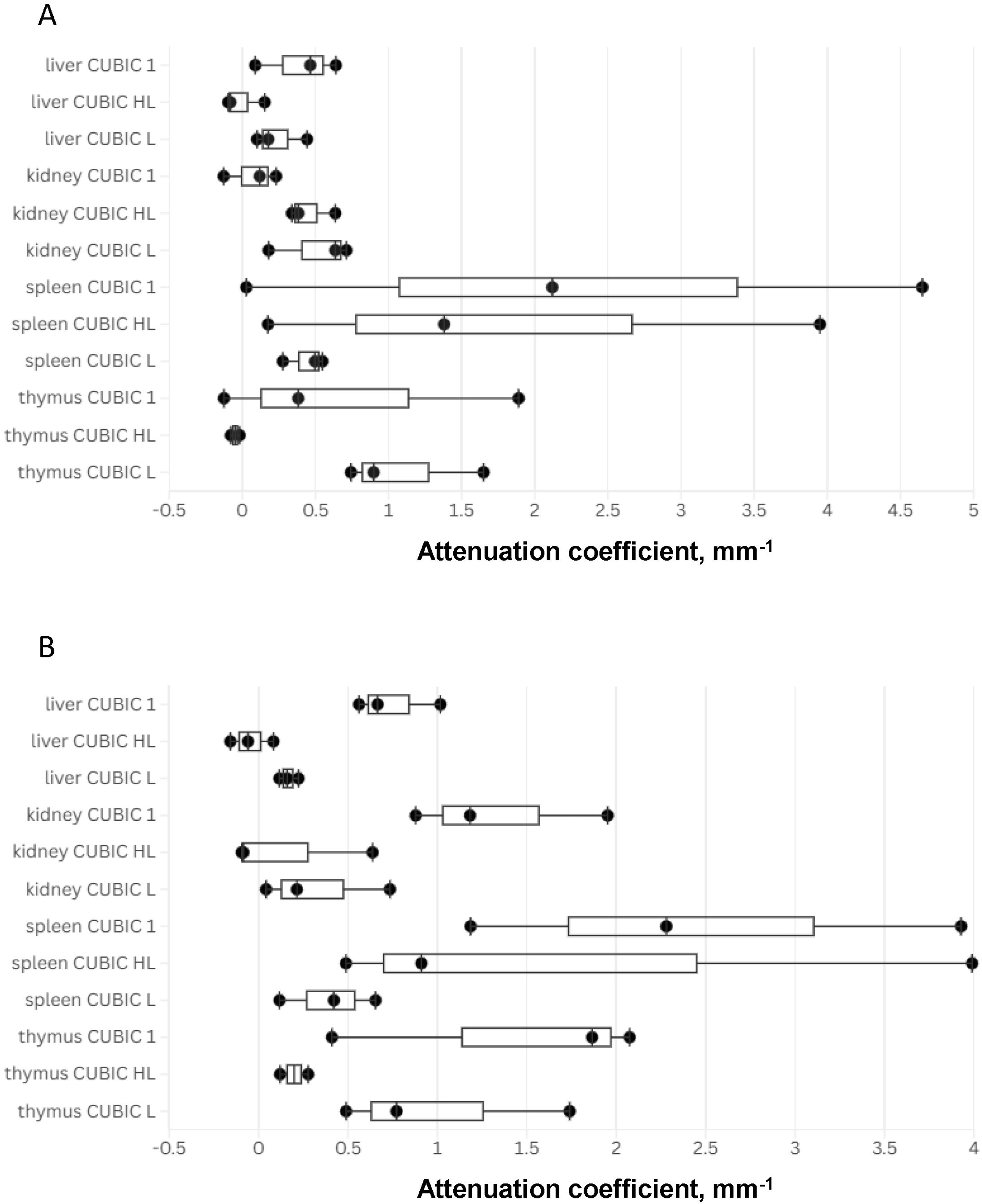
Fluorescence intensity attenuation coefficients of the global intensity profiles in the autofluorescence (A) and propidium iodide (B) channels for the tissues cleared by the three protocols. Boxplots (1st quartile, median, 3rd quartile) are shown for visualization only; all experimental points (one point per specimen) are displayed as filled circles. Very high attenuation coefficients (9.9 mm-1 for autofluorescence, 9.7 mm-1 for propidium iodide) for the non-cleared liver tissue (Fig. 8) are not shown on the plots for better visualization of the differences between the coefficients of the cleared specimens.

### Additional specimens

Furthermore, we applied CUBIC 1/CUBIC 2 and CUBIC L/CUBIC RA protocols to thymi from transgenic mice expressing mTurquoise and mCherry fluorescent proteins in the nuclei. In agreement with publications [3,10], fluorescence proteins signal was preserved after the both clearing protocols, permitting us to acquire 3D LSFM images (S2). However, in the CUBIC L/CUBIC RA-treated thymus, substantial fluorescence background occurred across the tissue in the mTurquoise channel, being most likely the autofluorescence that was rather bright in CUBIC L-treated “non-FP” specimens described above (Figs 3-7).

## Discussion

In the present work, we proposed a workflow for robust integral estimation of the tissue clearing and fluorescence staining quality, which is based on the depth-dependent attenuation of fluorescence in the tissue. The workflow uses the resulting three-dimensional fluorescence images themselves — rather than simple readouts (like specimen transmittance) — as a metric for assessing clearing quality; thus the workflow integrates multiple factors that influence the quality of the final LSFM images (which constitute the ultimate objective of tissue clearing and fluorescent staining).

Using this workflow, here we systematically compared clearing of various murine organs by the three protocols based on CUBIC reagents – 1) CUBIC 1/CUBIC 2, 2) CUBIC L/CUBIC RA, 3) CUBIC HL/CUBIC RA - with modifications (namely, nuclei staining was performed simultaneously with RI matching, and RI of CUBIC RA solution was adjusted to 1.45, to optimally match the requirements of Lightsheet.Z1 optics). Autofluorescence detected in the “GFP” channel permitted us to visualize tissue context in a label-free way, therefore we consider it beneficial when a channel is dedicated to autofluorescence only. Some differences in clearing efficiency and in PI nuclei staining occurred between the protocols. For instance, CUBIC L showed the lowest fluorescence attenuation coefficients among the tested protocols (for all organs except for thymi), indicating the most efficient clearing and PI staining. Thus, when an optimal clearing and staining protocol for a given tissue has not been established, then the CUBIC L/CUBIC RA protocol should be considered as a first option. This recommendation is based on its overall efficient tissue clearing, robust PI staining, and good preservation of fluorescent protein signals. However, when autofluorescence in “CFP” or “GFP” channels is undesirable (when a specific label emits in one of these channels), then CUBIC 1/CUBIC 2, yielding a dimmer autofluorescence, would be more suitable than CUBIC L/CUBIC RA. Furthermore, harsh CUBIC HL reagent is known to clear “difficult” tissues, such as human brain [10]; in our experiments, it was the most efficient for some tissues. However, CUBIC HL has disadvantages as well: chemical hazards due to its high pH, incompatibility with fluorescent proteins, and risk of dissolution of the specimens. In our opinion, CUBIC HL is worth trying if more “gentle” delipidation reagents such as CUBIC L or CUBIC 1 did not yield a good clearing (in the present study, that was the case for thymi). Moreover, when using CUBIC HL, one should carefully optimize duration of incubation, to find a balance between efficient delipidation and preservation of the specimen mechanical integrity.

Beyond transparency, a critical parameter of most tissue clearing protocols is the quality of specific fluorescence labelling (in the present study, nuclei labelling with PI). Staining of CUBIC-cleared tissues with PI has already been reported, as an additional three day-long step performed after delipidation [25]. Here, we demonstrated that PI staining can be performed simultaneously with refractive index matching in CUBIC RA or CUBIC 2 solutions, thereby reducing the overall duration of tissue clearing&staining protocols. Considering a long time typically required for each step of whole-mount specimen treatment, we think that combination of RI matching and fluorescent staining is worth to try also within other clearing+staining protocols. PI dissolved in CUBIC RA yielded a bright and uniform nuclei staining through the depth of the tissue; PI in CUBIC 2 resulted in a depth-attenuated, but still acceptable nuclei staining. That attenuation of PI signal could not be solely explained by lower tissue transparency, since in the same specimens, autofluorescence signal (dependent on transparency only) was less attenuated than that of PI. Therefore, other factor(s) have contributed to the PI signal attenuation; these factors could be a less efficient penetration of PI into tissues and/or its reduced affinity to DNA in CUBIC 2. Regardless to the exact physico-chemical mechanism, the practical conclusion is that PI staining in CUBIC 2 is acceptable, but less efficient than that in CUBIC RA.

CUBIC reagents permitted to clear a whole mouse [3], suggesting their capacity to clear all murine tissues. In our study, at least one of the three tested CUBIC protocols was efficient for each of the five tested tissues, including the heavily pigmented ones: liver, kidney and spleen, in agreement with the known capacity of CUBIC reagents to solubilize heme [10].

Nevertheless, few differences between the tested organs occurred in our experiments. For kidney and liver, all the three tested protocols were efficient, yielding rather low intensity attenuation coefficients (<0.6 mm^−1^, Fig 9A). For spleens, only CUBIC L/CUBIC RA appeared to be efficient, while two other protocols yielded irreproducible results (Fig 9). Thymi were efficiently cleared only by CUBIC HL, while CUBIC L and CUBIC 1 yielded wide sample-to-sample variations (Fig 9) and even inhomogeneity of transparency within the same specimen (Fig 6F). The most notable case was intestine; efficient clearing and PI staining was achieved by both CUBIC L/CUBIC RA and CUBIC 1/CUBIC 2, presumably thanks to thin walls and hollow shape of that organ. On the other hand, this anatomic organization has likely favored quick loss of mechanical integrity of intestine fragments in CUBIC HL. (As a side note, we did not calculate the global intensity profiles for intestine, because they would have depended on their anatomical features, such as wall thickness).

Thus, the observed differences between protocols and organs illustrate that a known efficiency of a clearing&staining protocol for a given tissue may only be extrapolated with great caution to other tissues or other clearing protocol variants. Our observations emphasize that several protocols are worth testing to achieve optimal clearing and staining of a tissue of interest.

One should note that the current version of our workflow is not an integrated, fully automated pipeline. It combines the steps performed in Imaris (with its GUI or batch processing tools), and Python-scripted steps. We consider it as a prototype which can be developed into a more automated pipeline in the future. Furthermore, the major limitation of our approach is its poor suitability to high throughput, because clearing and staining of whole organs and large tissue fragments, as well as LSFM image acquisition (including specimen mounting in the chamber) are time-consuming and difficult to automate. We realize that simpler readouts (namely, light transmittance) are more suitable for intermediate- or high-throughput experiments, such as screening of clearing reagents. The advantages of our approach are the following: it does not extrapolate simple readouts to the quality of volumetric fluorescence images; it is free of spatial sampling bias; it permits to evaluate contributions of transparency and specific fluorescent staining quality to the final fluorescence images. In conclusion, our workflow would be most useful for comparison of global performance of a few tissue clearing&fluorescence staining protocols or conditions, which can be precisely tailored to the specimens and fluorescent labels of interest.

## Acknowledgements

We acknowledge: Dr. Jose Morin Lantero for the custom Python script for calculations of the global fluorescence intensity profiles, as well as Dr. Ayele Denboba, Ms. Brigitte Krauth and Ms. Christiane Happe-Grammer for kindly providing freshly excised murine organs and advice on their handling.

