## Supplementary figures and images for "Volumetric fluorescence microscopy-based quantitative comparison of murine tissue clearing using CUBIC protocols"

### Fig S1

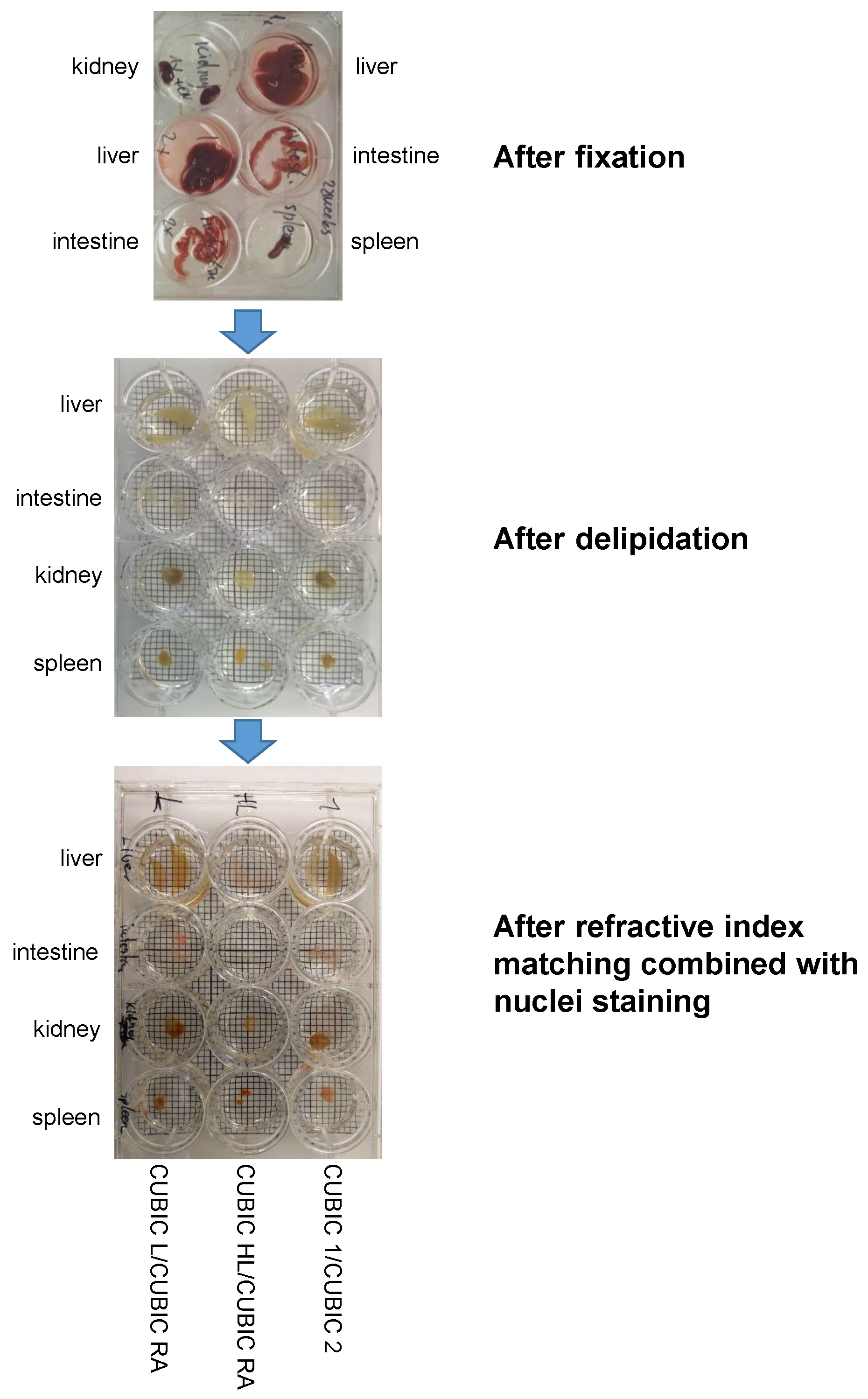

### Fig S2

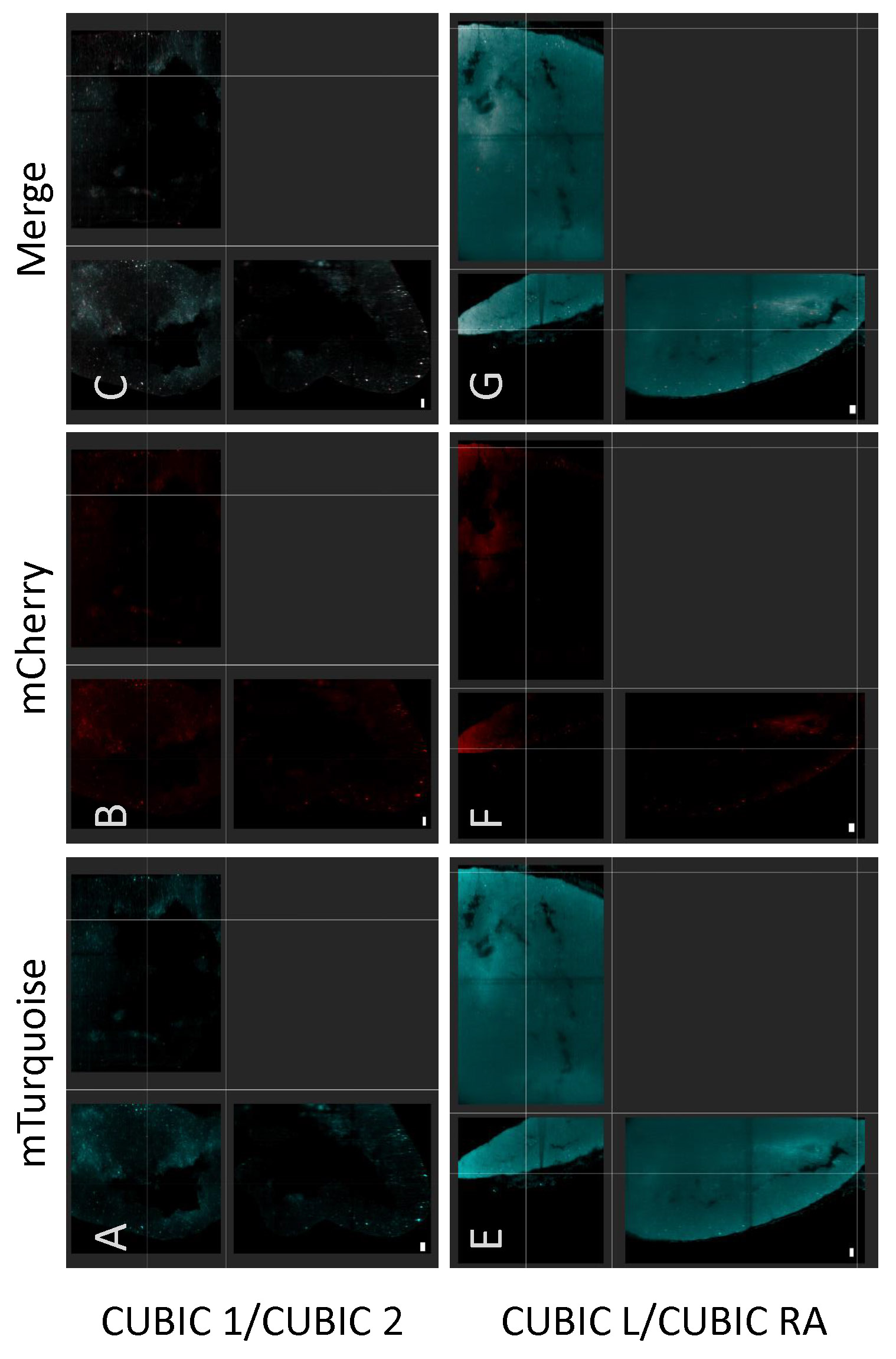

### Fig S3

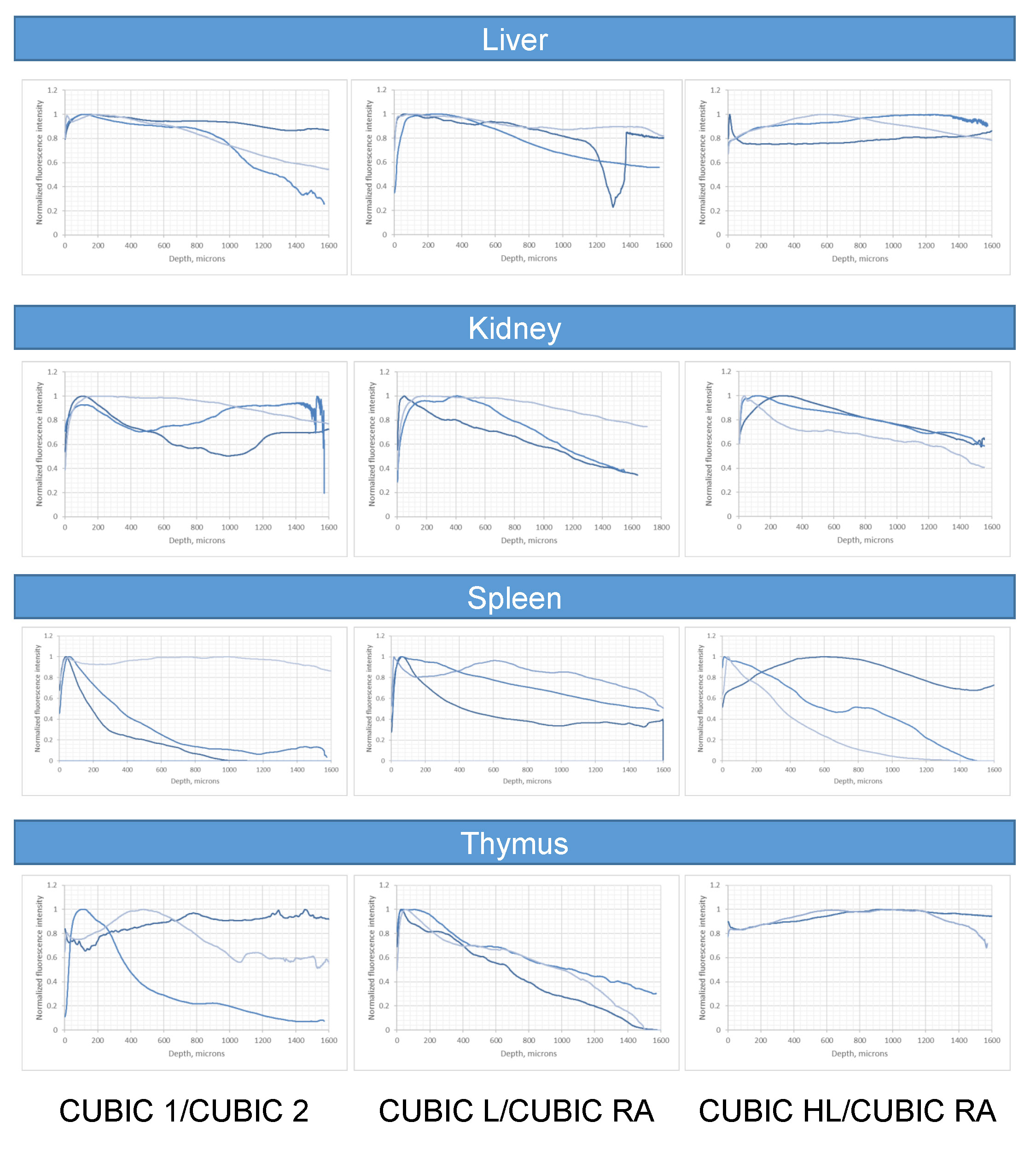

### Fig S4

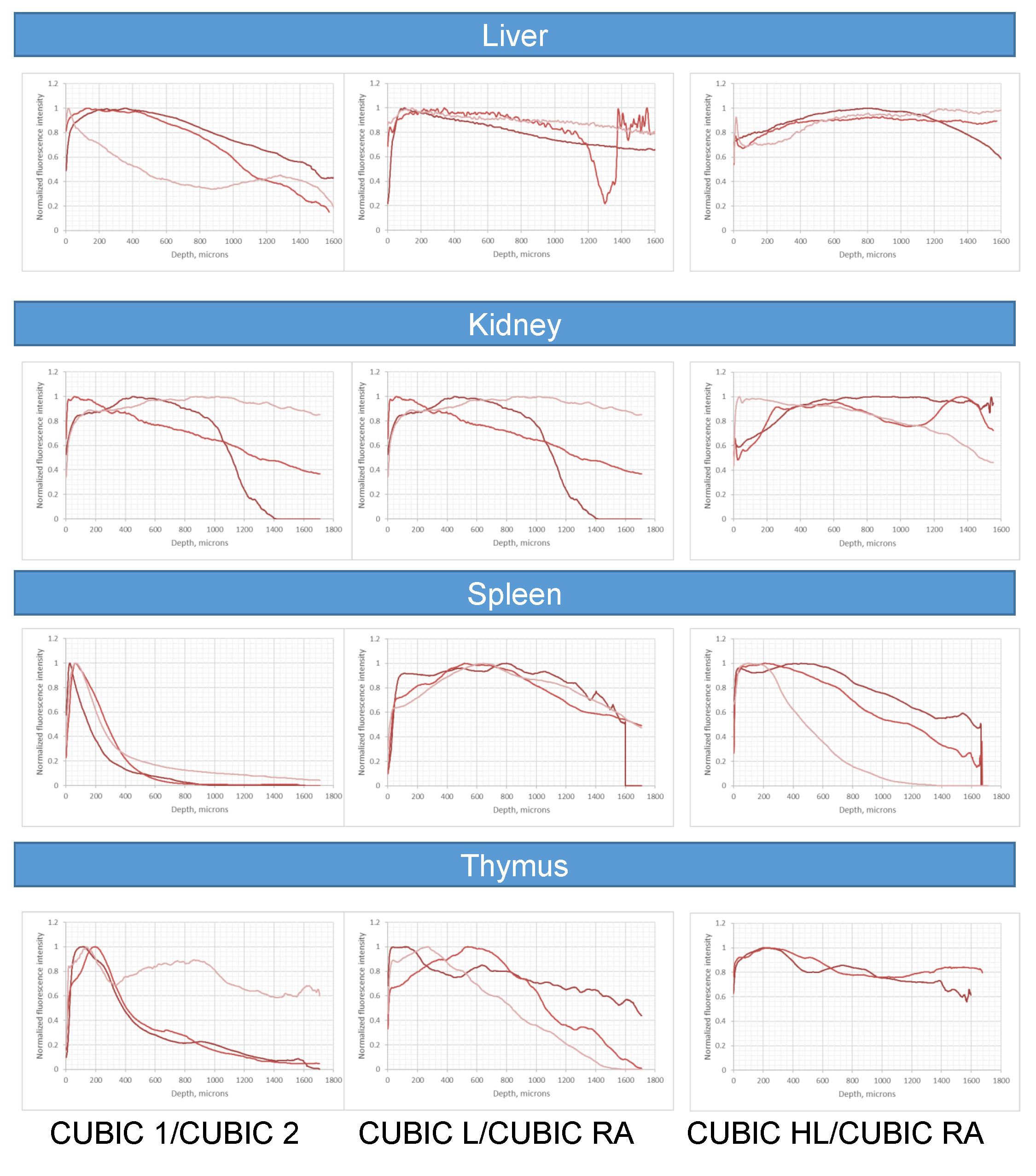
